# Escalating protein supersaturation underlies inclusion formation in muscle proteinopathies

**DOI:** 10.1101/762245

**Authors:** Prajwal Ciryam, Matthew Antalek, Fernando Cid, Gian Gaetano Tartaglia, Christopher M. Dobson, Anne-Katrin Guttsches, Britta Eggers, Matthias Vorgerd, Katrin Marcus, Rudolf A. Kley, Richard I. Morimoto, Michele Vendruscolo, Conrad Weihl

## Abstract

Abundant, aggregation prone or “supersaturated” proteins are a feature of neurodegeneration. Whether the principle of supersaturation can similarly explain the widespread aggregation that occurs in non-neuronal protein conformational disorders and underlies pathogenic protein aggregate formation is not established. To test this prediction we analyzed proteomic datasets of biopsies from genetic and acquired protein aggregate myopathy (PAM) patients by quantifying the changes in composition, concentration and aggregation propensity of proteins in the fibers containing inclusions and those surrounding them. We found that similar to neurodegeneration, a supersaturated subproteome of aggregate prone proteins is present in skeletal muscle from healthy patients. This subproteome escalates in degree of supersaturation as proteomic samples are taken more proximal to the pathologic inclusion, eventually exceeding its solubility limits and aggregating. While most supersaturated proteins decrease or maintain steady abundance across healthy fibers and inclusion containing fibers, supersaturated proteins within the aggregate subproteome rise in abundance, suggesting they escape normal regulation. We show in the context of a human conformational disorder that the level of supersaturation of a metastable subproteome helps to explain widespread aggregation and correlates with the histopathological state of the tissue.

**Significance:** Increasing evidence implicates the phenomenon of protein supersaturation with the selective vulnerability of specific cells to protein misfolding disorders. Quantitative studies of this phenomenon, however, have only been possible post mortem in the case of neurodegenerative diseases. To overcome this limitation, we study here protein aggregate myopathies (PAMs), for which we were able to carry out systematic single fiber proteomic studies on patient-derived samples. We found not only that proteins associated with PAM inclusions are highly supersaturated in muscle but also that their supersaturation levels increases further in affected fibers. These results provide a clear illustration of how an escalation in supersaturation leads protein inclusions in vulnerable cells.

## Introduction

The presence of protein aggregates is a hallmark of many age-related degenerative disorders (1, 2). These aggregates are characteristic of neurodegenerative diseases, but are also features of disorders outside of the central nervous system, including protein aggregate myopathies (PAMs) (3). One unifying hypothesis relating to the pathogenesis of these proteinopathies is the age-related disruption of the protein homeostasis system (1, 2). For example, mutations in aggregation-prone proteins or changes in the cellular environment promote protein misfolding and subsequent aggregation in affected tissues (4, 5). These aggregation events lead to further progressive impairment in protein surveillance and degradation pathways, causing further aggregation of other aggregation-prone proteins.

To rationalize these observations, we recently proposed that protein aggregation is a widespread phenomenon associated with the intrinsic supersaturation state of the proteome (6, 7). As proteins are supersaturated when their cellular concentration exceeds their solubility, we have developed a metric combining a sequence-based prediction of aggregation propensity and estimates of protein concentration for transcriptomic and proteomic data to estimate the supersaturation levels of thousands of human proteins (7). With this approach, we have reported that proteins found in inclusions in Alzheimer’s disease (AD), Parkinson’s disease (PD), and amyotrophic lateral sclerosis (ALS) have high supersaturation scores even in control tissues (7, 8). We have also similarly shown that the proteins that aggregate in aging *C. elegans* are supersaturated (7).

Remarkably, the enrichment for supersaturated proteins in neurodegenerative pathways is apparent even when estimating supersaturation levels from average abundances across a wide variety of tissues. However, the tissue selectivity of many protein conformational disorders suggests that the risk of misfolding may depend in part on the specific proteomic context. A limitation of previous studies on supersaturation is the absence of this context, because of the intractability of obtaining living brain tissue from patients with neurodegenerative disease (5, 9). Because muscle is easily biopsied, the PAMs offer a means to determine how proteinopathies can remodel specific tissues, and whether changes in supersaturation in this context help to explain disease progression and pathology. In these degenerative muscle disorders, protein accumulates into inclusion bodies in affected myofibers (3, 10). In some cases these inclusions contain the same proteins associated with neurodegenerative diseases, such as TDP-43 and SQSTM1 (10).

Most hereditary PAMs are due to a dominantly inherited missense mutations in specific proteins resulting in their destabilization and subsequent aggregation (3). By contrast, sporadic inclusion body myositis (IBM) is an acquired PAM with no clear genetic etiology manifesting exclusively in patients over 45 years of age (11). Two types of pathologic structures exist in PAMs, inclusion bodies, which are often immunoreactive for the mutated protein in hereditary disease, and rimmed vacuoles (RVs), which are pathologic structures found in affected myofibers and contain aggregated proteins in association with degradative debris such as ubiquitin and autophago-lysosomal proteins (11). In these diseases, as in other neurodegenerative disorders, the aggregation of particular disease-related proteins initiates a series of pathological changes that ultimately give rise to aggregation. In the present study, we use quantitative proteomic data from human patient tissues to test the hypothesis that superstation of a metastable subproteome explains protein inclusions in PAMs. Moreover, we explore how the aggregate subproteome changes between healthy cells, diseased cells and inclusion bearing cells.

## Results

### IBM-associated proteins are supersaturated in healthy tissues

We previously performed laser microdissection to collect areas of single fibers from muscle biopsies of 18 patients with IBM (12). These samples were taken from normal healthy fibers, or in the case of IBM-affected muscles, from affected RV-containing fibers and adjacent normal appearing fibers. We then analyzed these samples by mass spectrometry using label-free spectral count-based relative protein quantification (see Methods). For the study presented here, healthy control and IBM proteomic datasets were generated from healthy control myofiber regions (HCs), unaffected myofiber regions from IBM patients (disease controls, DCs), non-vacuole containing sarcoplasmic regions of affected fibers (AFs), and myofiber regions containing rimmed vacuoles (RVs) (**Figure 1A**).

**Figure 1.**
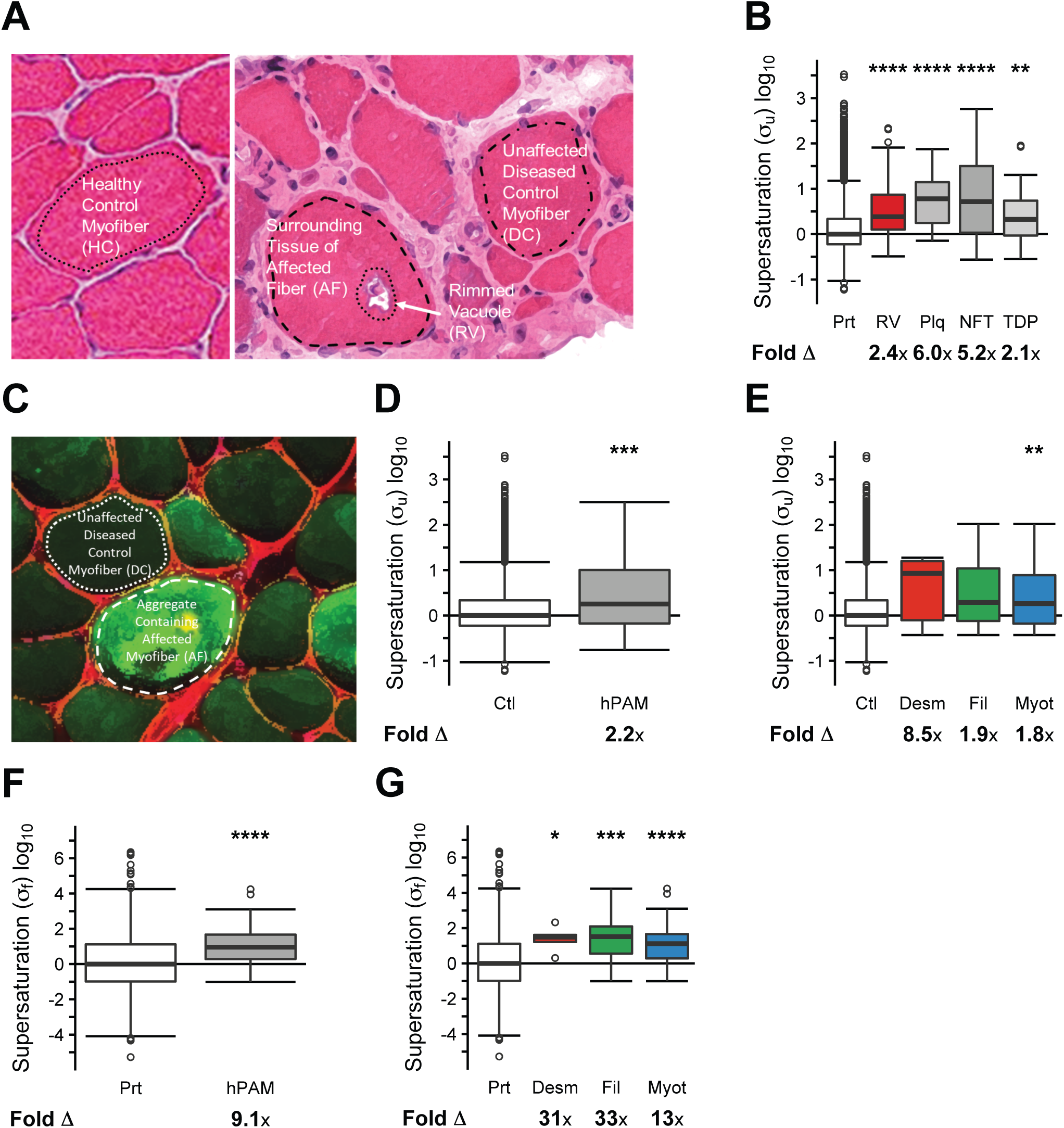
Proteins in rimmed vacuoles from protein aggregation myopathies are supersaturated. Representative images of: **(a)** healthy control myofibers (HC), control unaffected myofibers in diseased samples (DC), surrounding tissues of affected fibers (AF), and rimmed vacuoles (RV) from human subjects with inclusion body myositis, and **(c)** DC and AF samples from human subjects with myotilin mutations. Outlines represent areas for LMD. In **(c)**, prior to LMD, muscle was immunostained with an antibody directed to myotilin (green) to identify aggregate containing fibers (AF). **(b, d, e)** Comparison of the unfolded supersaturation scores (σ_u_) of the proteome (Prt) (N=15954) and (**b)** proteins enriched in RVs (RV) (N=50), amyloid plaques (Plq) (N=26), neurofibrillary tangles (NFT) (N=76), proteins found in TDP-43 inclusions (TDP) (N=32); **(d)** proteins enriched in affected fibers from any of three protein aggregation myopathies (hPAM) (N=50); and **(e)** proteins enriched in affected fibers from individual protein aggregation myopathies involving desmin (Desm) (N=6), filamin (Fil) (N=16), and myotilin (Myot) (N=46) mutations. **(f,g)** Comparison of the folded supersaturation scores (σ_f_) for the proteome (Prt) (N=1605) and **(f)** the proteins enriched in affected fibers from any of three protein aggregation myopathies (hPAM) (N=46) and **(g)** the proteins enriched in affected fibers from individual protein aggregation myopathies involving desmin (Desm) (N=5), filamin (Fil) (N=15), and myotilin (Myot) (N=43) mutations. The fold change (Δ) represents the fold difference in the median σ_u_ or σ_f_ scores between each inclusion type and the proteome. The median σ_u_ or σ_f_ supersaturation score for the proteome is normalized to 0. Boxes range from the 25^th^ percentile to 75^th^ percentile, while whiskers extend to maximum and minimum data points up to 1.5x interquartile range above and below the limits of the boxes. Remaining outliers are plotted as open circles. Statistical significance estimated by one-tailed Wilcoxon/Mann-Whitney test with Holm-Bonferroni correction. *p < 0.05, **p < 0.01, ***p < 0.001, ****p < 0.0001.

Comparison of these proteomic datasets enabled us to identify a set of proteins statistically enriched within RVs, as compared to DCs. This list of 53 RV-enriched proteins includes 17 proteins previously identified to accumulate in IBM tissue (**Dataset S1**). We next asked whether these proteins share similar biophysical features despite their different sequences, structures and functions. We had previously estimated supersaturation of a protein as the product of its predicted aggregation propensity (given by the Zyggregator score (*Z*_*agg*_) which correlates negatively with its solubility) and its expression level, either based on mRNA levels from microarray data or proteomic analysis (7).

We thus compared the supersaturation levels of RV-enriched proteins to those of co-aggregators within amyloid plaques (13), neurofibrillary tangles (14), and TDP-43 inclusions (8) (**Dataset S2**). As a gross approximation of the supersaturation level for a given protein, we used mRNA levels averaged over dozens of different human tissues unaffected by misfolding disease (**Dataset S4**) and aggregation propensities predicted from the primary sequences for the unfolded states of proteins (*Z*_*agg*_) (**Dataset S3**) termed the unfolded supersaturation score (σ_u_) (**Dataset S2**) (4). While this approach does not benefit from tissue specificity, it was previously shown that even this average estimate demonstrated elevated supersaturation scores for proteins associated with aggregation and cellular pathways implicated in neurodegenerative disorders and enables us to compare directly inclusions from muscle and the central nervous system (7).

We found that proteins enriched in RVs have elevated supersaturation scores (σ_u_) in control tissues (RV, median fold change (Δ): 2.4x, p=1.4•10^−6^). This was similar to proteins observed to co-aggregate (co-aggregators) with plaques (median Δ: 6.0x, p=4.5•10^−8^) and neurofibrillary tangles (median Δ: 5.2x, p=1.3•10^−13^) in AD, and TDP-43 (median Δ: 2.1x, p=1.8•10^−3^) in ALS, respectively (**Figure 1B, Dataset S7**). The elevated supersaturation score for RV enriched proteins was also present when we calculated tissue-specific supersaturation scores 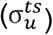 using the subset of the cross-tissue microarray expression database that included skeletal muscle expression (RV: median Δ: 2.1x, p=2.2•10^−6^) (**SI Appendix, Figure S1**; **Dataset S7**). Comprehensive statistical results are shown in **Dataset S12**.

### hPAM-associated proteins are supersaturated in healthy tissues

To determine whether the phenomenon of supersaturation observed for IBM-associated proteins (**Figure 1B**), an acquired PAM, is also observed for proteins associated with hereditary PAMs (hPAM), we extended our studies to proteomic datasets from laser microdissected myofibers of muscle biopsies of patients with three different genetically defined hPAMs (10 patients with *DES* mutations, 7 patients with *FLNC* mutations and 17 patients with *MYOT* mutations) (15-17). Samples were taken from affected aggregate-containing fibers (AF) or adjacent normal appearing disease control fibers (DC) (**Figure 1C**). We then identified statistically enriched proteins within the aggregate-containing fibers, as compared to unaffected disease control fibers (**Dataset S2**). The σ_u_ is similarly elevated for the proteins enriched in hPAM aggregate fibers (AF) (median Δ: 2.2, p=6.9•10^−4^) (**Figure 1D, Dataset S7**). We note, however, that sample size limitations led to statistically insignificant results for two of the three individual hPAMs (desminopathy median Δ: 8.5x, p=9.8•10^−2^; filaminopathy median Δ: 1.9x, p=8.3•10^−2^, myotilinopathy median Δ: 1.8x, p=6.7•10^−3^) (**Figure 1E**). We then calculated 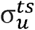 and found the increased supersaturation of proteins in aggregate-containing tissue is significant in this context (hPAM: median Δ: 4.5x, p=1.2•10^−8^; desminopathy median Δ: 11x, p=2.5•10^−2^; filaminopathy median Δ: 5.6x, p=1.4•10^−3^, myotillinopathy median Δ: 3.9x, p=7.5•10^−7^) (**SI Appendix, Figure S1**; **Dataset S7**). In addition, we estimated the significance of the relative increase in supersaturation of the 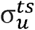 scores relative to the σ_u_ scores (p < 1•10^−6^).

Particular to our proteomic datasets, we can evaluate the degree of supersaturation within the context of the skeletal muscle proteome rather than abundances from mRNA levels in public databases. Thus we asked whether aggregate-enriched proteins in hPAMs were supersaturated, based on their abundance in background of a healthy muscle proteome (**Dataset S6)**. To do this, we combined protein abundances derived from healthy control muscle with a version of the Zyggregator score that weights residue-level aggregation propensities based on predictions of the relative burial of proteins after folding 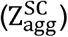, as described previously (18) (**Dataset S3**). This estimate is termed folded supersaturation score (σ_f_) as compared with the previous estimate of the unfolded supersaturation score (σ_u_) (**Dataset S9)**. To directly compare these two estimates (σ_u_ and σ_f_), we calculated the σ_f_ of proteins enriched in hPAM aggregate-containing fibers (median Δ 9.1x, p=5.3•10^−5^) (**Figure 1F** compared with **Figure 1D)**. This elevated supersaturation score among proteins enriched in hPAM aggregate-containing fibers relative to HC is a result of both abundances and aggregation propensities higher than those of the proteome 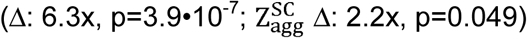 (**SI Appendix, Figure S2A-B, Datasets S3 and S9**). Similarly, we found elevated σ_f_ when considering proteins enriched in aggregate-containing tissue specific to each hPAM (**Figure 1G, Dataset S2)** (desminopathy median Δ: 31x, p=1.4•10^−2^; filaminopathy median Δ: 33x, p=8.8•10^−4^; myotillinopathy median Δ: 13x, p=5.7•10^−5^).

In unaffected diseased control myofibers (DC), we found that the σ_f_ of proteins enriched in aggregate-containing myofibers (AF) were elevated relative to the proteome (desminopathy median Δ: 16x, p=1.2•10^−2^; filaminopathy median Δ: 4.6x, p=5.7•10^−2^; myotilinopathy median Δ: 4.4x, p=2.7•10^−3^) (**Figure 2A, D, G**). In comparison, the σ_f_ of these proteins in affected fibers (AF) were higher (desminopathy median Δ: 1700x, p=4.3•10^−5^; filaminopathy median Δ: 48x, p=4.2•10^−5^; myotilinopathy median Δ: 33x, p=3.6•10^−8^) (**Figure 2B, E, H**). To confirm the robustness of these results, we introduced varying amounts of random noise into our data, and found that the results are robust even when noise of at least 5 times the magnitude of the signal, and in many cases as much as 20x the magnitude of the signal, is introduced (**SI Appendix, Figures S3-S4**).

**Figure 2.**
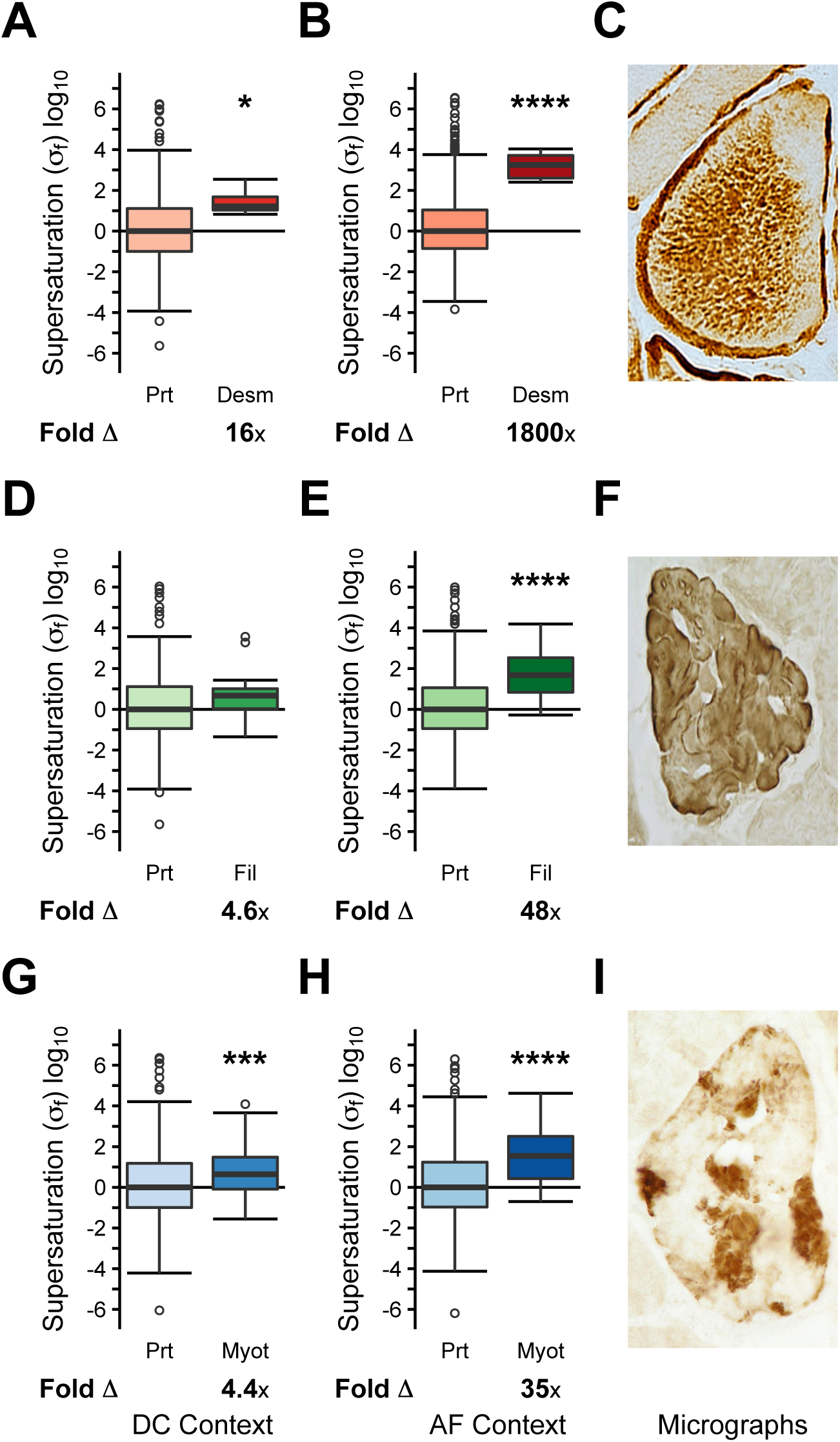
Protein supersaturation in hereditary protein aggregate myopathies. Comparison of folded supersaturation scores (σ_f_) between the proteome and proteins enriched in aggregate-containing myofibers (AF) for: **(a,b,c)** desminopathy (DC: Prt N=611, Agg N=6; AF: Prt N=1387, Agg N=6), **(d,e,f)** filaminopathy (DC: Prt N=333, Agg N=16; AF: Prt N=507, Agg N=16), and **(g,h,i)** myotillinopathy (DC: Prt N=680, Agg N=46; AF: Prt N=742, Agg N=46). Box plots and statistical tests as in **Figure 1**. *p < 0.05, **p < 0.01, ***p < 0.001, ****p < 0.0001.

### Escalating supersaturation in IBM

By utilizing our IBM proteomic datasets, we had segmented data starting from healthy controls (HC) and continuing to unaffected fibers in affected patients (DC), areas from affected fibers surrounding the RV (AF), and the RV itself (RV) (**Figure 1A, Figure 3A-E, Datasets S6 and S9**). These data allowed us to determine how the σ_f_ of the proteins that are enriched in RVs transition from healthy fibers to aggregate-containing fibers. To obtain this result, we calculated σ_f_ based on protein abundances for each of these contexts. Even in healthy controls, the σ_f_ of RV-enriched proteins is higher in the muscle context than what we found in the cross-tissue transcriptional analysis for unfolded supersaturation (σ_u_) (median Δ: 7.3x, p=9.6•10^−4^) (comparison p < 2.0•10^−5^) or with the skeletal muscle unfolded supersaturation score 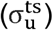 (median Δ: 2.1x, p=2.2•10^−6^) (**Figure 3A** compared with **Figure 1B** and **SI Appendix, Figure S1**). In healthy control (HC) fibers, this result is driven by the higher median aggregation propensity of RV-enriched proteins rather than an increase in abundance (abundance Δ: 1.9x, 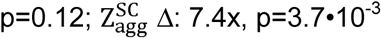) (**SI Appendix, Figure S2C-D**).

**Figure 3.**
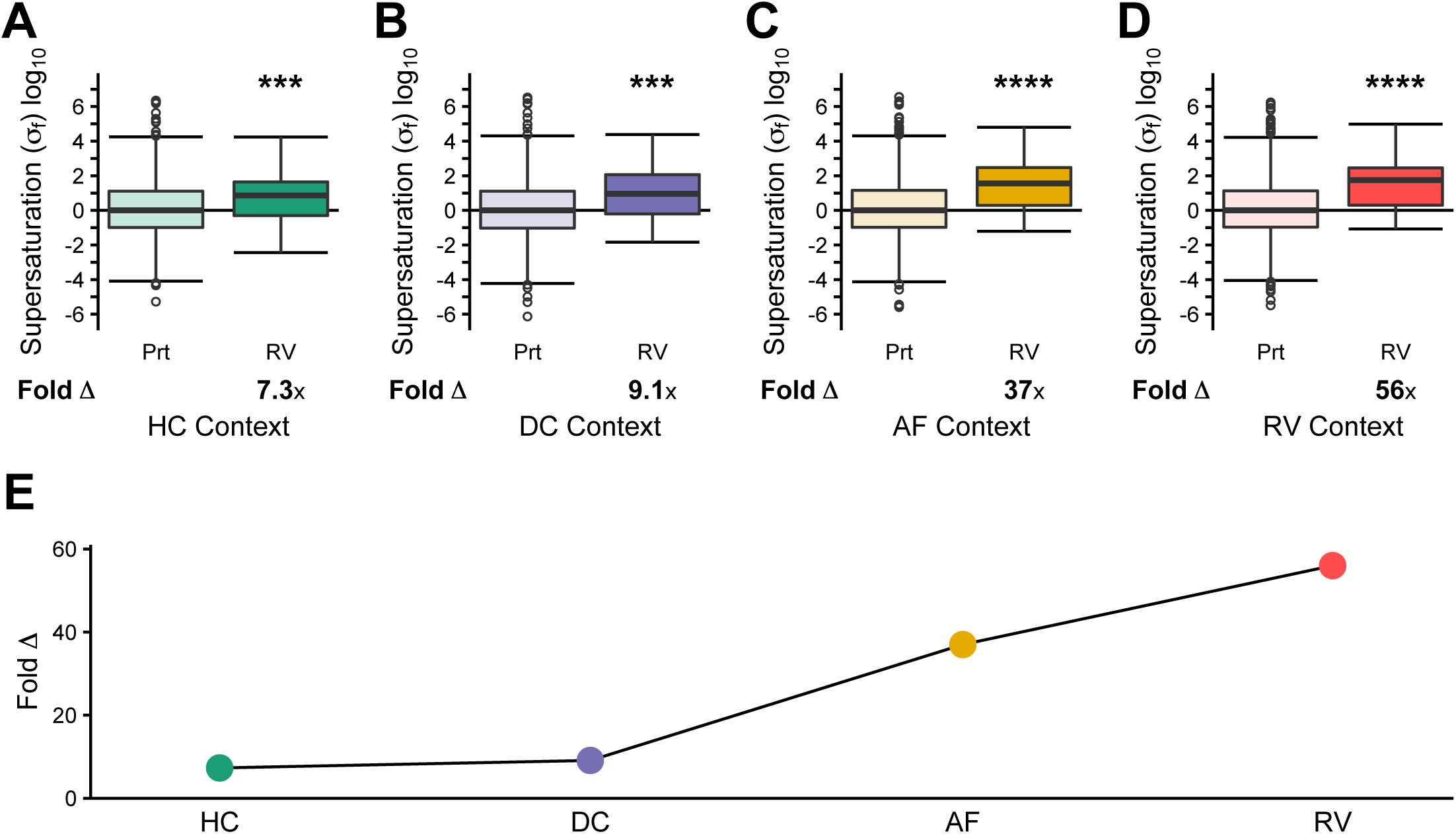
Escalating supersaturation in inclusion body myositis. Comparison of folded supersaturation scores (σ_f_) for the proteome (Prt) and proteins enriched in rimmed vacuoles (RV) relative to diseased control myofibers. **(a)** Healthy control myofiber (HC) (Prt N=1605, RV N=47), **(b)** control myofibers unaffected in diseased samples (DC) (Prt N=1988, RV N=52), **(c)** aggregate-containing affected myofibers (AF) (Prt N=2396, RV N=52), and **(d)** rimmed vacuoles (RV) (Prt N=2104, RV N=52). **(e)** Comparison of the fold difference in median σ_f_ between RV and Prt. Box plots and statistical tests as in **Figure 1**. *p < 0.05, ***p < 0.001, ****p < 0.0001.

We found that σ_f_ increases with the physical proximity to the RVs (DC median Δ: 9.1x, p=1.2•10^−4^; AF median Δ: 37x, p=1.2•10^−9^; RV median Δ: 56x, p=4.3•10^−10^) (**Figure 3B-E**). These results are robust against high levels of noise (**SI Appendix, Figure S5**-**S6**). In order to determine whether this rise in supersaturation was statistically significant, we performed a simulation of 1,000,000 trials in which we randomly selected 53 proteins to determine how frequently we could achieve a similar pattern of results by chance. The combination of elevated supersaturation scores in each context with rising median Δ has an estimated of p=0.011 (see Supplementary Methods). Given that this analysis included proteins expressed in some contexts but not in others (e.g. present in disease fibers but not healthy control fibers), we confirmed that the results were qualitatively similar when considering only a limited set of proteins expressed in all four contexts and had associated Zyggregator scores available (**SI Appendix, Figure S7, Dataset S9**). To further validate these results, we performed these analyses utilizing aggregation predictions from the unfolded state with *Z*_*agg*_ and TANGO (19), which similarly demonstrated a significant escalation in supersaturation (**SI Appendix, Figure S8-S9, Dataset S10).**

Like proteins enriched in RVs, proteins enriched in hPAM aggregate-containing fibers also exhibit an escalating σ_f_ in the sporadic disease context (**SI Appendix, Figure S10**). The escalation in σ_f_ is specific for proteins that accumulate in PAMs since proteins that co-aggregate with amyloid plaques (**SI Appendix, Figure S11A-E**) and neurofibrillary tangles (**SI Appendix, Figure S11F-J**) in AD do not exhibit escalating σ_f_ in IBM muscle tissues.

### RV proteins escape the downregulation of supersaturated proteins

We recently reported a transcriptional suppression of supersaturated proteins and pathways in Alzheimer’s disease (20). We therefore asked whether a similar phenomenon takes place at the transcriptional and translational level in IBM. To do so, we determined the proteins differentially expressed in affected fibers (AF) relative to healthy controls in IBM (HC) (**Dataset S11**). We found, across independent patient datasets, that 52 proteins are decreased and only one protein, desmin, is increased in affected fibers. Those proteins that are decreased in the surrounding fibers tend to have higher σ_f_ in healthy controls relative to the rest of the proteome (median Δ: 3.8x, p=9.8•10^−5^) (**Figure 4A**). There are 830 proteins (Prt) in our dataset for which we had 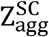 in HC context and abundance values across all four contexts, and of these, only 48 (5.8%) are decreased in abundance in affected fibers. By contrast, of the top 5% most supersaturated proteins in this subset (N=41) (Top σ_f_), seven (17%) are decreased in affected fibers (enrichment p=0.013) (**Figure 4B**). As further validation of this phenomenon, we used RNA sequencing data from healthy muscle and IBM muscle to identify the transcripts of proteins that were downregulated in IBM tissue (21). The downregulated transcripts correspond to proteins whose supersaturation scores tend to be elevated in healthy controls (median Δ: 2.6x, p=3.3•10^−3^) (**SI Appendix, Figure S12A**). There are 1366 transcripts for proteins in this dataset for which we were able to calculate σ_f_ in HC context. Of the top 5% most supersaturated proteins in this subset (N=68) (Top σ_f_), 15 (22%) are decreased in expression in affected fibers versus 157 (11%) for the proteome (Prt) as a whole (enrichment p=0.016) (**SI Appendix, Figure S12B**).

**Figure 4.**
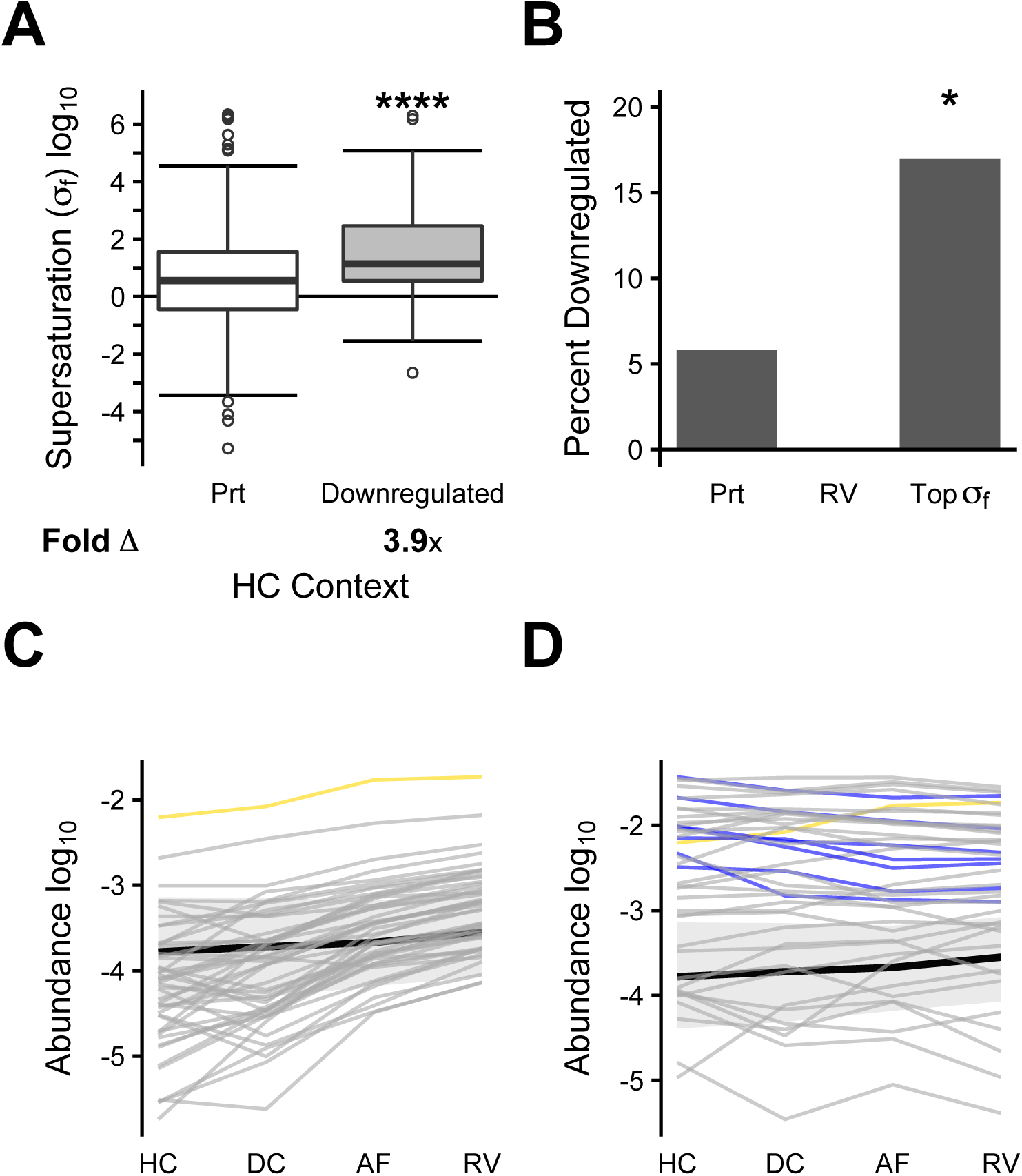
Protein supersaturation is associated with downregulation. In this analysis, only proteins that are detected in HC, DC, AF, and RV, and for which there are defined σ_f_ scores in HC are included. **(a)** Folded supersaturation scores (σ_f_) for the proteome (Prt) (N=830) and proteins downregulated from HC to AF (N=50). Box plots and statistical tests as in **Figure 1. (b)** Percentage of proteins downregulated in the proteome (Prt) (48/830), proteins enriched in rimmed vacuoles (RV) (0/47), and top 5% most supersaturated proteins (based on HC context) (Top σ_f_) (7/41). Significance estimated by the Fisher Exact test, with Holm-Bonferroni correction. **(c)** Protein abundances in HC, DC, AF, and RV are plotted for the 47 proteins enriched in RVs included in the subset analyzed in this figure. Desmin is highlighted in yellow, the only RV-enriched protein whose abundance is increased between HC and AF. **(d)** Protein abundances in HC, DC, AF, and RV are plotted for the 47 proteins with the highest supersaturation scores (top 5%). Desmin again is highlighted in yellow, the only highly supersaturated protein with rising abundance. Those proteins that are significantly downregulated between HC and AF are highlighted in blue. In **(c)** and **(d)**, the background black line and grey bar represent median and 25^th^-75^th^ percentile range for the 830 proteins in the proteome in this subset. *p < 0.05, ****p < 0.0001.

The RV-enriched proteins (RV) are an exception, as none of them is downregulated at the protein (**Figure 4B**) or transcript (**SI Appendix, Figure S12B**) level. The individual abundance trajectories of these RV-enriched proteins trend towards rising abundances, although only one of the proteins, desmin, significantly rises in abundance between HC and AF (**Figure 4C**). By contrast, among the top 5% most supersaturated proteins, there is a trend towards declining abundances, with a disproportionate number of proteins decreasing in abundance significantly in this group (**Figure 4D**). These results suggest that supersaturated proteins are typically downregulated to control their abundance, but when they fail to be downregulated, they escape regulation and are more likely to deposit into inclusions.

Only one protein is among the top 5% most supersaturated proteins, an RV-enriched protein, and increases significantly in abundance. This protein is desmin, mutations in which are associated with desminopathy and which is found to be enriched in aggregate-associated tissues in myotilinopathy, filaminopathy and IBM. Desmin represents the clearest example of an escape protein and is also highly associated with protein misfolding in muscle tissue.

## Conclusions

By using protein abundance data from proteomic datasets derived from human biopsy specimens, we identified a metastable, supersaturated subproteome in muscle tissue from protein aggregate myopathies. These data are consistent with our previous studies that explored the phenomenon of supersaturation in proteinopathies associated with neuronal inclusions such as AD and ALS (7, 8). In contrast to those previous studies, the current analysis evaluated supersaturation using the protein abundances from the affected tissues rather than estimated abundances averaged across tissues obtained from publicly available datasets. Thus, skeletal muscle offers a unique opportunity to explore how the proteome remodels in the face of aggregation-related disease, and the ways in which this can be rationalized by the physicochemical characteristics of solubility and expression. Our analysis identified PAM-specific subproteomes of metastable proteins that behaved similarly in the context of supersaturation analysis. The proteins that are found in RVs and inclusions in disease have elevated supersaturation scores even in healthy tissue, suggesting that there is an intrinsic risk for aggregation even in the normally expressed proteome.

Our ability to analyze samples taken from unaffected and affected myofibers within the same patient enabled demonstration that the degree of supersaturation escalates from normal myofibers, to unaffected diseased myofibers, to aggregate containing myofibers. In the case of IBM, the quality of the data enabled us to show an escalating supersaturation to the RV from surrounding tissue within the same fiber. To our knowledge, a confirmatory demonstration that an aggregate prone subproteome increases in abundance from unaffected to affected cells in a vulnerable tissue from human biopsy specimens has never been performed.

In both IBM and the three hereditary myopathies (hPAM) we studied, aggregate-associated proteins have elevated supersaturation scores in the context of healthy tissue (HC). In addition, affected fibers (AF) have higher relative supersaturation scores than unaffected fibers in patients known to have the disease (DC), both in the sporadic and hereditary cases. There are likely multiple factors contributing to the progressively rising abundance (and hence, supersaturation level) of proteins that deposit in RVs. In part, this may reflect a failure in proteostatic machinery, as has been shown in a variety of conformational disorders (7). Our results suggest that there may also be a failure to suppress the expression of some highly supersaturated proteins, given that those proteins that deposit in RVs run counter to a trend of declining abundance for supersaturated proteins (**Figure 5**). That this signal is apparent at the transcriptional level, as well, favors a role for dysregulation of abundance, as opposed merely to a failure in the function of molecular chaperones or degradation machinery. Our findings suggest that affected fibers have the capacity to downregulate their supersaturated proteome, and that this occurs at least in part at the transcriptional level. These data are consistent with our previous study in AD, in which downregulated proteins are similarly supersaturated relative to the proteome (20). These results suggest that there may be a mechanism in IBM by which supersaturated proteins are preferentially downregulated to maintain their solubility. Indeed, the abundances of the top 5% most supersaturated proteins in skeletal muscle remained stable or decreased as they approached the RV. In contrast, supersaturated RV enriched proteins tended to increase in abundance. These analyses identified one abundant protein, desmin, which was enriched within the RV and appears to escape the downregulation common to other highly abundant proteins.

**Figure 5.**
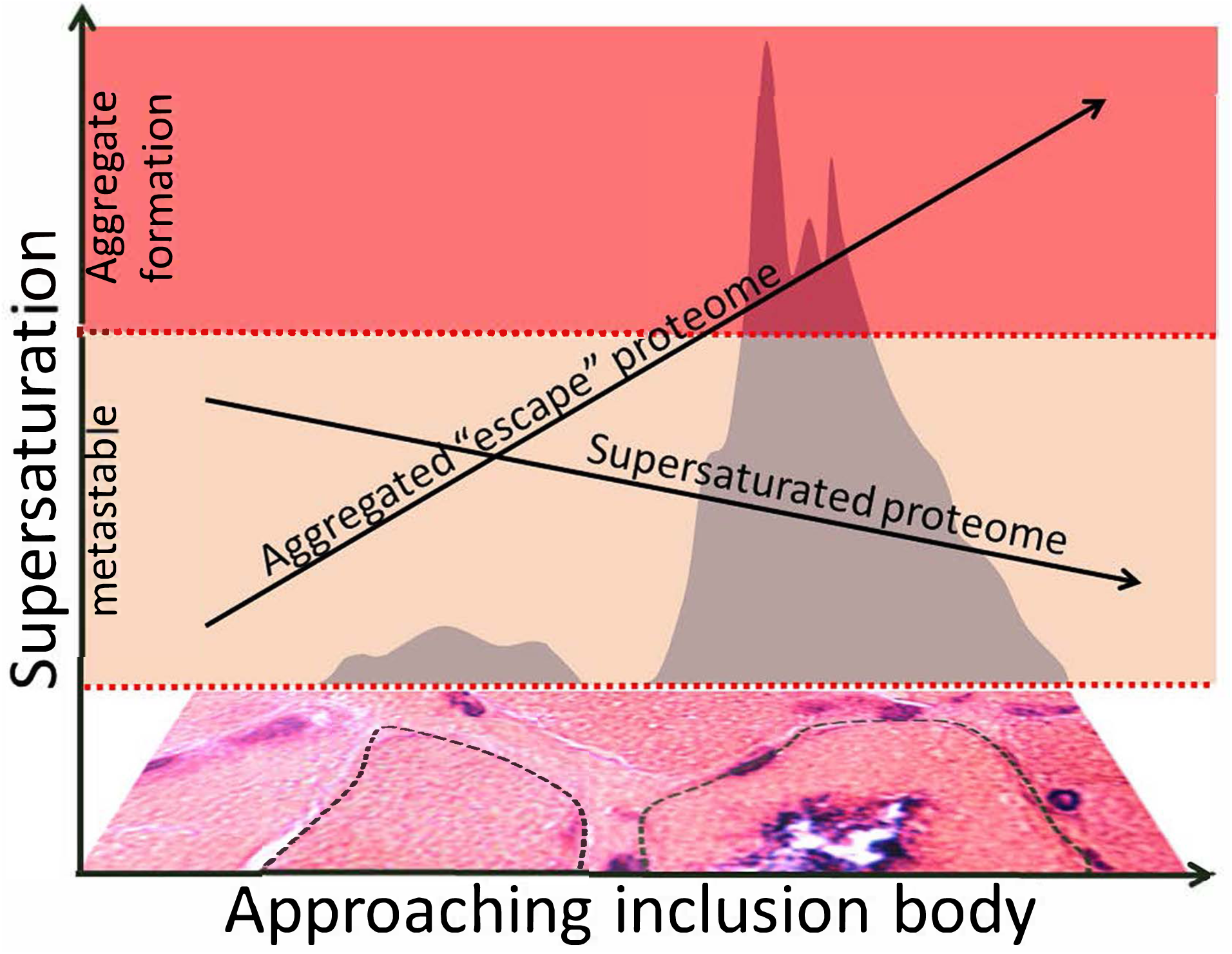
Escalating supersaturation of an aggregate prone subproteome puts affected fibers at risk of inclusion formation in inclusion body myositis. 1) Supersaturation of the aggregate proteome increases to the point of aggregate formation at muscle inclusion bodies (gray regions). 2) The most highly supersaturated proteins decrease in abundance upon approaching the inclusion body. In contrast, the abundance of the aggregate/RV enriched supersaturated proteome increases and escapes downregulation.

The pathogenic mechanism associated with supersaturation suggests that a protein or subset of proteins reach a critical concentration that exceeds their solubility resulting in protein aggregation (7). Therapeutic approaches aimed at buffering this metastable proteome may be effective at reducing the degree of supersaturation. The present study identifies a common subset of abundant and aggregation prone proteins from >50 well characterized patients with PAMs. These proteins include amyloidogenic proteins such as gelsolin and TDP-43 that are not mutated in genetic PAMs but are mutated in hereditary amyloidosis and ALS respectively. These findings suggest that therapies aimed at their reduction may also be effective at restoring protein homeostasis in PAMs. Our observation that desmin is both supersaturated in healthy control tissue and rises in abundance with the muscle’s pathological progression makes it a potential target for such intervention. Finally the degree of supersaturation of a subset of proteins may serve as proxy for the proteostatic state of muscle. One could envision using the concentration of the aggregate proteome as a biomarker in future therapies focused on PAMs. Taken together, our results indicate that the presence of supersaturated proteins represents a persistent challenge for the protein homeostasis system, and that failures in regulating the aggregation of these proteins leads to the formation of inclusions in a wide range of diseases, including neurodegenerative disorders and protein aggregate myopathies.

## Methods

### Datasets

The datasets used in this work and the proteins in each of them are described in **Datasets S1 and S2**. Description of data analysis and statistical measures are in supplemental methods.

### Laser microdissection (LMD) and sample processing

Patients provided informed consent. Study protocols were approved by the local ethics committee (reg. number 3882-10) at Ruhr-University Bochum, Bochum, Germany. For each patient 250,000 µm^2^ of HC, DC, AF or RV tissue was collected by LMD (LMD 6500, Leica Microsystems, Wetzlar, Germany). Sample lysis and digestion were carried out as previously described (16). Briefly, samples were lysed with formic acid (98-100%) for 30 minutes at room temperature (RT), followed by a sonication step for 5 min (RK31, BANDELIN electronic, Berlin, Germany). Samples were kept frozen at −80 °C until digestion.

Prior to digestion the formic acid was removed and the collected samples were digested in 50 mM ammonium bicarbonate at pH 7.8. Samples were reduced and alkylated by adding dithiothreitol and iodoacetamide. Trypsin (Serva) was added to a final concentration of 1 µg. Digestion was carried out overnight at 37 °C and stopped by adding TFA to acidify the sample. Samples were purified using OMIX C18 Tips (Varian, Agilent Technologies, Böblingen, Germany) completely dried vacuum and again solved in 63 µl 0.1% TFA, as described in (16).

### Mass spectrometry

16 µl per sample were analysed by nano-liquid chromatography tandem mass spectrometry (nanoLC-ESI-MS/MS). The nano high performance liquid chromatography (HPLC) analysis was performed on an UltiMate 3000 RSLC nano LC system (Dionex, Idstein, Germany) as described in (15). Peptides were separated with a flow rate of 400 nl/min using a solvent gradient from 4% to 40% B-solvent for 95 min. Washing of the column was performed for 5 min with 95% B-solvent and was then returned to 4% B-solvent. The HPLC system was online-coupled to the nano electrospray ionization (ESI) source of an Orbitrap elite mass spectrometer (Thermo Fisher Scientific, Germany). Mass spectrometric measurements were performed as previously described (12).

## Acknowledgements

Research was supported by the National Institute of Health grant number R21NS101588-01A1, R01AR068797 and K24AR073317 to CW and a generous gift from Catalyze for a Cure.

