## Appendix for "Escalating protein supersaturation underlies inclusion formation in muscle proteinopathies"

**Supplemental Methods**

**Data analysis.** Raw files were converted into the Mascot generic format (MGF) format using Proteome Discoverer 1.4 (Thermo Fischer Scientific, Germany). MGF files were searched against a combined database containing the Swiss-Prot part of the UniProt Knowledgebase (UniProtKB) (1) or homo sapiens (release 2014/05/28, 20265 curated entries). For the generation of shuffled decoy entries DecoyDatabaseBuilder was used (2). Identifications were performed by Mascot 2.5 (Matrixscience Ltd, (3)) with a peptide mass tolerance of 10 ppm, fragment mass tolerance of 0.5 Da, one allowed missed cleavage and carbamidomethylation (C), oxidation (M) as well as phosphorylation (S, T, Y) as variable modifications. Label free relative quantification by spectral counting was performed as described in (4).

***Calculation of protein abundance***. We previously reported abundances as spectral counts normalized by the total number of spectral counts in a given sample (4-7). Here, we performed an additional normalization step to account for the fact that longer proteins will generate more peptides in mass spectrometry than smaller proteins of the same abundance (8). Akin to the normalized spectral abundance factor, we divided normalized spectral counts in our data sets by the protein length. We then divided these values by the sum of all such normalized values in a given sample. We then averaged these normalized protein abundances across replicates and $\log_{10}$-transformed these values to arrive at a final abundance value.

***Calculation of gene expression from microarray data.*** Microarray data was obtained from BioGPS pre-processed using gcrma as previously described. For cross-tissue analysis, cell ine and malignant tissue expression levels were excluded. Transcript identifiers were converted to UniProt IDs, with cases of ambiguous conversion or absence of reviewed UniProt IDs excluded from analysis. Values ≤ 0 were excluded. Expression levels were then $\log_{10}$-transformed then averaged across all values for a given UniProt ID. A similar procedure was done for the skeletal muscle analysis, but limiting it to the two arrays of skeletal muscle data.

***Calculation of gene expression from RNA sequencing data.*** Processed RNA sequencing data was obtained, with expression levels reported in FKPM (GEO Datasets GSE102138) (9). Any values ≤ 0 were excluded. Identifiers were converted to reviewed UniProt IDs, with ambiguous conversions excluded from further analysis. In cases in which one multiple identifiers mapped to a single UniProt IDs, these FKPM values were averaged. The values were then $\log_{10}$-transformed. Significantly upregulated and downregulated transcripts were identified based on the reported q-values. In cases in which there were multiple q-values associated with a given UniProt ID, the largest q-value was used. The q-values reported were two-tailed, which we converted to one-tailed q-values for the purpose of our analysis. We used a threshold of significance of $p < 0.05$.

***Calculation of protein aggregation propensity***. For the human proteome set, we calculated the $Z_{agg}$, $Z_{agg}^{SC}$ and TANGO scores as previously described (10, 11). For TANGO, we set the parameters at pH = 7.4, T = 310K, and ionic strength = 0.1M. The supersaturation score $\sigma$ is calculated as the sum

$$\sigma = C + Z \qquad (S1)$$

where C is the $\log_{10}$ of the concentration and $Z$ is aggregation propensity score; the concentrations are derived from the protein abundance levels. In each dataset, values were recentered such that the median $\sigma$ score for each database was 0.

***Identification of proteins enriched in disease-associated inclusions.*** In order to determine vacuole-enriched proteins in the IBM data set, we compared abundance values in the RV dataset to those in the DC dataset. For this analysis, we only included proteins that had a non-zero abundance in both the DC and RV datasets, which constituted a total of 1302 proteins. For these proteins, we performed a one-tailed paired t-test. We then used the Benjamini-Hochberg method to calculate q-values for each of these proteins, using as a threshold of significant q < 0.05, for a False Discovery Rate of 5%.

***Gaussian noise generation***. We performed noise testing to evaluate the robustness of our results for the comparison of supersaturation scores among the IBM data sets, as well as the hPAM data sets. We defined one hundred noise levels on the basis of the standard deviation of a series of Gaussian distributions with mean of 0. The range of standard deviations was $\log_{10}(1.1)$ to $\log_{10}(10.1)$. At each noise level $l$, we performed 100 trials $t$, in which we drew a random number $n_{l,t,p}$ from that the noise level distribution for each of the $p$ proteins in the database. The noise-introduced supersaturation score $\sigma_{p,l,t}$ was defined as

$$\sigma_{p,l,t} = \sigma_p + n_{l,t,p} \qquad (S2)$$

For trial $t$ of noise level $l$, the set $S_{l,t}$ of noise values is

$$S_{l,t} = \{n_{l,t,1}, n_{l,t,2}, \dots, n_{l,t,p}\} \qquad (S3)$$

The set $m_{l,t}$ of linear magnitudes of noise for trial $t$ of noise level $l$ is

$$m_{l,t} = \left\{10^{\ln\lceil n_{l,t,1} \rceil}, 10^{\ln\lceil n_{l,t,2} \rceil}, \dots, 10^{\ln\lceil n_{l,t,p} \rceil}\right\} \qquad (S4)$$

For noise level $l$, the set $M_l$ of median noise values for its constituent trials is

$$M_l = \{median(m_{l,1}), median(m_{l,2}), \ldots, median(m_{l,100})\} \qquad (S5)$$

In each Gaussian noise plot, the values plotted on the x-axis were the median of $M_l$ with error bars representing the standard error of the mean as calculated using default settings in the Python package SciPy.

***Gaussian noise significance testing.*** For each trial at each noise level, we determined the sets of noise-modified $\sigma$ scores for the data sets under consideration. A one-tailed Wilcoxon/Mann-Whitney U test was performed for each of these trials, with multiple hypothesis correction performed based on the same families used for the original analysis, with one difference. At each noise level, the median of the p-values for the 100 trials was plotted with error bars representing the standard error of the mean as calculated using default settings in the Python package SciPy. We performed a one-sided one-sample t-test using the distribution of p-values for a given trial to test the null hypothesis that the mean of these p-values is not significantly less than 0.05. For those cases in which we could not reject the null hypothesis, we plotted the points in grey; otherwise, we plotted the points in color.

***Gaussian noise fold change testing.*** For each trial at each noise level, we determined the sets of noise-modified $\sigma$ scores for the data sets under consideration. The linear difference $d_{l,t}$ between the medians of the supersaturation scores of the control set $C_{l,t}$ and experiment set $E_{l,t}$ being tested at noise level $l$ and trial $t$ is

$$d_{l,t} = 10^{median(E_{l,t}) - median(C_{l,t})} \qquad (S9)$$

At noise level l, we plotted the median of set $\{d_{l,1}, d_{l,2}, \ldots, d_{l,100}\}$ with error bars representing the standard error of the mean as calculated using default settings in the Python package SciPy. We performed a one-sided one-sample t-test using the distribution of fold change values for a given trial to test the null hypothesis that the mean of these fold changes is not significantly greater than 1. For those cases in which we could not reject the null hypothesis, we plotted the points in grey; otherwise, we plotted the points in color.

**Overlap analysis.** In **Figure 4B** and **SI Appendix, S12B**, the Fisher exact test is used to calculate enrichment of data sets for particular categories of proteins.

**Statistical significance of escalating supersaturation.** To test the significance of our observations of rising supersaturation (**Figure 3**, **SI Appendix, Figures S7-11**) we used a simulation. The null hypothesis was that it would arise by chance that 1) the median $\Delta > 0$ for a set of proteins of interest in each context and 2) median $\Delta$ of those proteins would rise successively from HC to DC to AF to RV contexts. To test this, we performed the following procedure $K$ times, where $K$ =1,000,000. For each trial $k$, we randomly selected $N$ proteins from the proteome (where $N$ is equal to the number of proteins of interest, for instance 53 in the case of RV-enriched proteins or 51 in the case of hPAM-enriched proteins). When selecting $N$, we used the total number of proteins meeting a particular criterion, even if a smaller number of those proteins was actually present in the original dataset. For these $N$ proteins, $D$ is the set of median $\Delta$ compared to the proteome for each of the four contexts:

$$D \equiv \{med\Delta_{HC}, med\Delta_{DC}, med\Delta_{AF}, med\Delta_{RV}\} \qquad (S10)$$

If the supersaturation rose successively at each from HC to DC to AF to RV, and median Δ > 0 in each context, we assigned a score $E_k$ of one; otherwise, we assigned a score $E_k$ of zero. We then summed this score over the 1,000,000 trials.

$$D = \{med\Delta_{HC}, med\Delta_{DC}, med\Delta_{AF}, med\Delta_{RV}\} \qquad (S11)$$

$$E_k = \begin{cases} 1, & if \min(D) > 0 \; and \; med\Delta_{RV} > med\Delta_{AF} > med\Delta_{DC} > med\Delta_{HC} \\ 0, & otherwise \end{cases} \qquad (S12)$$

We estimated the significance of the escalation in supersaturation as follows:

$$\mathrm{E} = \{\mathrm{E}_1, \mathrm{E}_2 \ldots, \mathrm{E}_\mathrm{K}\} \qquad (S13)$$

$$p = \sum_{k=1}^{K} \frac{E_k}{K} \qquad (S14)$$

$$p = \sum_{k=1}^{K} \frac{E_k}{K} \qquad (S15)$$

In order to test the isolated contribution of escalating median Δ, we removed the constraint of median Δ > 0, and calculated a score $E_k^r$:

$$E_k^r = \begin{cases} 1, & if med\Delta_{RV} > med\Delta_{AF} > med\Delta_{DC} > med\Delta_{HC} \\ 0, & otherwise \end{cases} \qquad (S16)$$

$$p = \sum_{k=1}^{K} \frac{E_k}{K} \qquad (S17)$$

We considered all cases analyzed by our original constraints on family for the purpose of multiple hypothesis correction and all cases analyzed by the relaxed criteria a separate family. Multiple hypothesis correction was performed using the Holm-Bonferroni method. P-values for both constraints are reported in **Dataset S12**.

**Statistical significance of comparative median Δ.** To test the significance of differences in median Δ between different contexts (**Figures 1-3**), we used a simulation. The null hypothesis was that the difference in median $\Delta$ ($\Delta_\Delta$), of at least the magnitude reported would arise by chance. The reported difference in median $\Delta$ we refer to as $\Delta_\Delta^0$. To test this, we performed the following procedure $K$ times, where $K$ =1,000,000. For each trial $k$, we randomly selected $N$ proteins from the proteome by the same procedure as above for escalating supersaturation. For these $N$ proteins, we calculated the median $\Delta$ in contexts $C_1$ and $C_2$. Note that we performed this analysis in a one-tailed fashion.

$S_{\Delta_\Delta} = \{\Delta_\Delta^1, \dots, \Delta_\Delta^K\}\Delta_\Delta = \{\Delta_\Delta^1, \dots, \Delta_\Delta^K\} \quad (S18)$, where

$$\Delta_\Delta^k = med\Delta_2^k - med\Delta_1^k \qquad (S19)$$

We assigned a score $E_k$ to each trial and from all the trials together derived a p-value, as follows:

$$E_k = \begin{cases} 1, & \Delta_\Delta^k > \Delta_\Delta^0 \\ 0, & otherwise \end{cases} \qquad (S20)$$

$$p = \sum_{k=1}^{K} \frac{E_k}{K} \quad (S21)$$

We considered all cases analyzed in this fashion as a single family. Multiple hypothesis correction was performed using the Holm-Bonferroni method. P-values are reported in **Dataset S12**.

***Multiple hypothesis correction.*** In order to perform adequate multiple hypothesis correction while avoiding increasing Type II error by overcorrecting, it was necessary to group our results into a series of families on which multiple hypothesis correction would be performed meaningfully. We used the following principles to help divide the analyses in these studies into a set of coherent families. Except when they were being compared directly, hPAM and IBM data sets were considered part of separate families. IBM families were organized cross data subsets (that is, HC, DC, AF, and RV included in the same family). hPAM families were organized in three families: 1) HC, 2) DC, and 3) AF. This was organized in this way because there were multiple individual hPAMs, but analyses for the composite group of hPAM aggregate-enriched proteins could only be performed logically on the HC dataset as the other data sets were disease-specific. Analyses using $\sigma_u$ were considered distinct from analyses using $\sigma_f$. All $\sigma_u$ analyses were considered as part of a single family. Among IBM data sets, we performed a series of analyses in which we compared $\sigma_f$ levels between the proteome and particular subsets of proteins (RV-enriched, hPAM-enriched, plaque-enriched, NFT-enriched) across the four IBM data sets (HC, DC, AF, RV). We considered analyses involving each of these subsets as separate families. **Dataset S12** shows a summary of all statistical tests performed in this analysis, and groups those tests by their respective families.

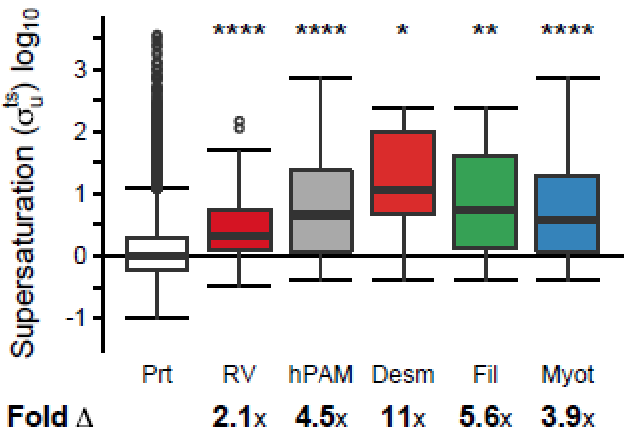


**Figure S1. Unfolded skeletal muscle specific supersaturation for aggregated proteins from proteins enriched in rimmed vacuoles and hereditary protein aggregate myopathies.** Comparison of skeletal muscle specific supersaturation scores ($\sigma_u^{ts}$) between the proteome and proteins enriched in rimmed vacuoles (RV) calculated using expression levels from microarray data obtained from skeletal muscle (Prt N=15944, RV N=50, hPAM N=49, desminopathy N=6, filaminopathy N=16, myotillinopathy N=45). Box plots and statistical tests as in **Figure 1**. *p < 0.05, **p < 0.01, ****p < 0.0001.

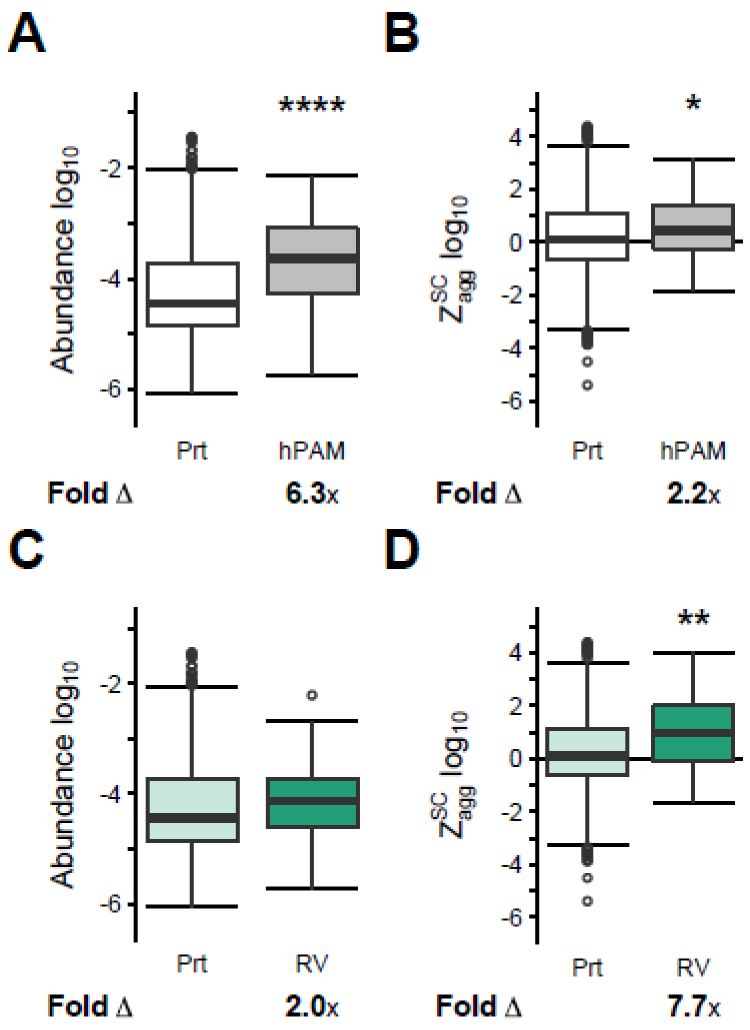


**Figure S2. Both abundance and aggregation propensity contribute to the elevated supersaturation of aggregation-prone proteins.** Comparison of the HC proteome (Prt) (N=1605) to (**a, b**) proteins enriched in affected fibers from any of three protein aggregation myopathies (hPAM) (N=46), or (**c**, **d**) proteins enriched in rimmed vacuoles (RV) (N=47). Results are provided in terms of: (**a**, **c**) protein abundance values estimated by mass spectrometry from healthy control myofibers, or (**b**, **d**) aggregation propensity scores (structurally corrected Zyggregator scores). Box plots and statistical tests as in **Figure 1**. *p < 0.05, **p < 0.01, ****p < 0.0001.

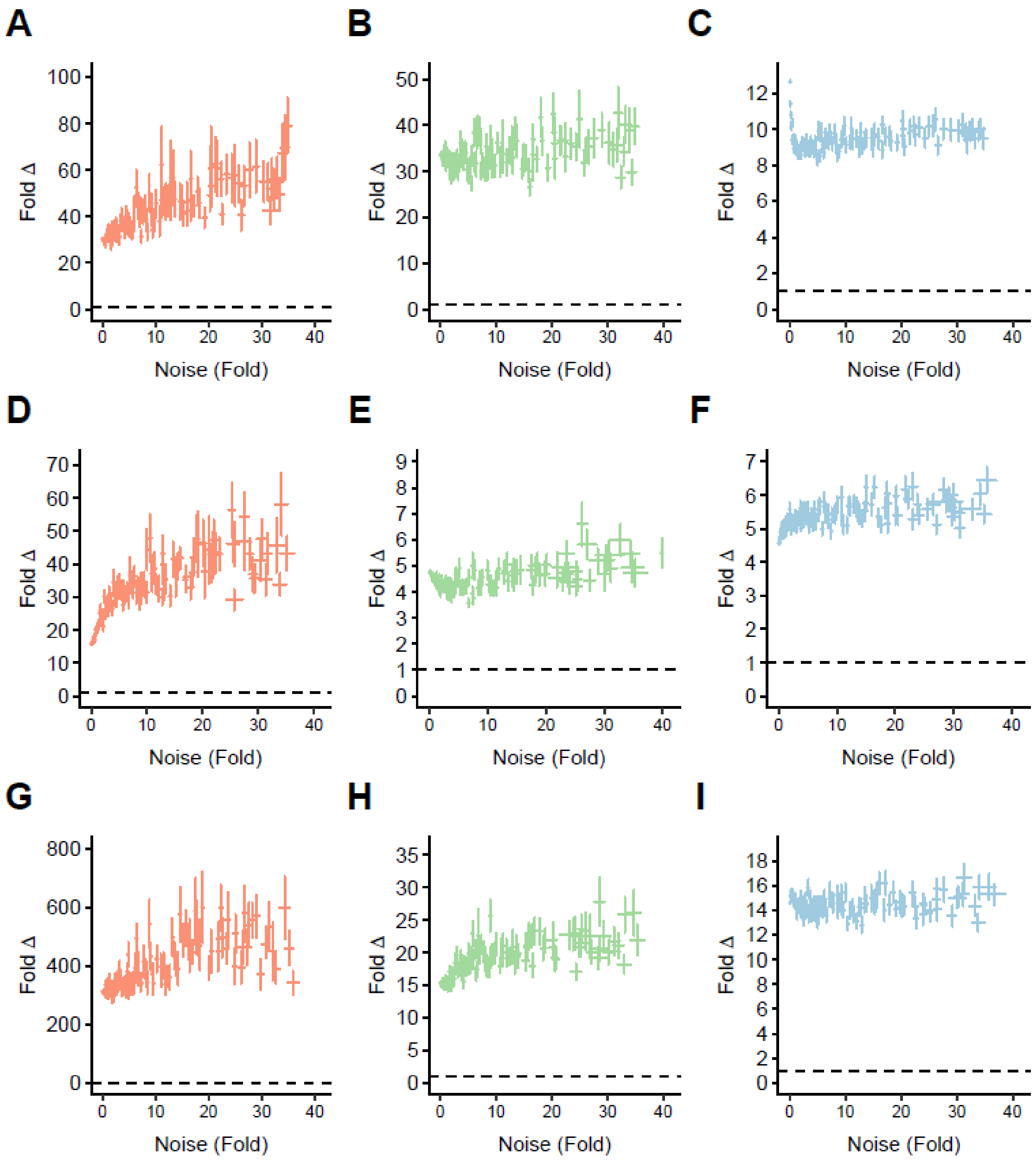


**Figure S3. Fold changes for supersaturation estimates for aggregation-prone proteins in individual hereditary protein myopathies are robust against random noise.** Random noise from increasingly wide Gaussian distributions was introduced into the protein supersaturation scores for the proteome and proteins enriched in affected fibers from hereditary myopathies, as shown in **Figure 2**. Points plotted are the mean ± S.E.M. of median fold difference between aggregate and proteome from 100 trials at each noise level based on: healthy control (**a-c**), disease control (**d-f**), or affected fiber context (**g-i**) for desminopathy (**a**, **d**, **g**), filaminopathy (**b**, **e**, **h**), and myotillinopathy (**c**, **f**, **i**), respectively. A one-tailed one-sample Student's t-test was performed at each noise level to determine whether median fold differences were significantly greater than 1 **(d-f)**. Colored points represent significant results. Dashed line marks median fold difference of 1.

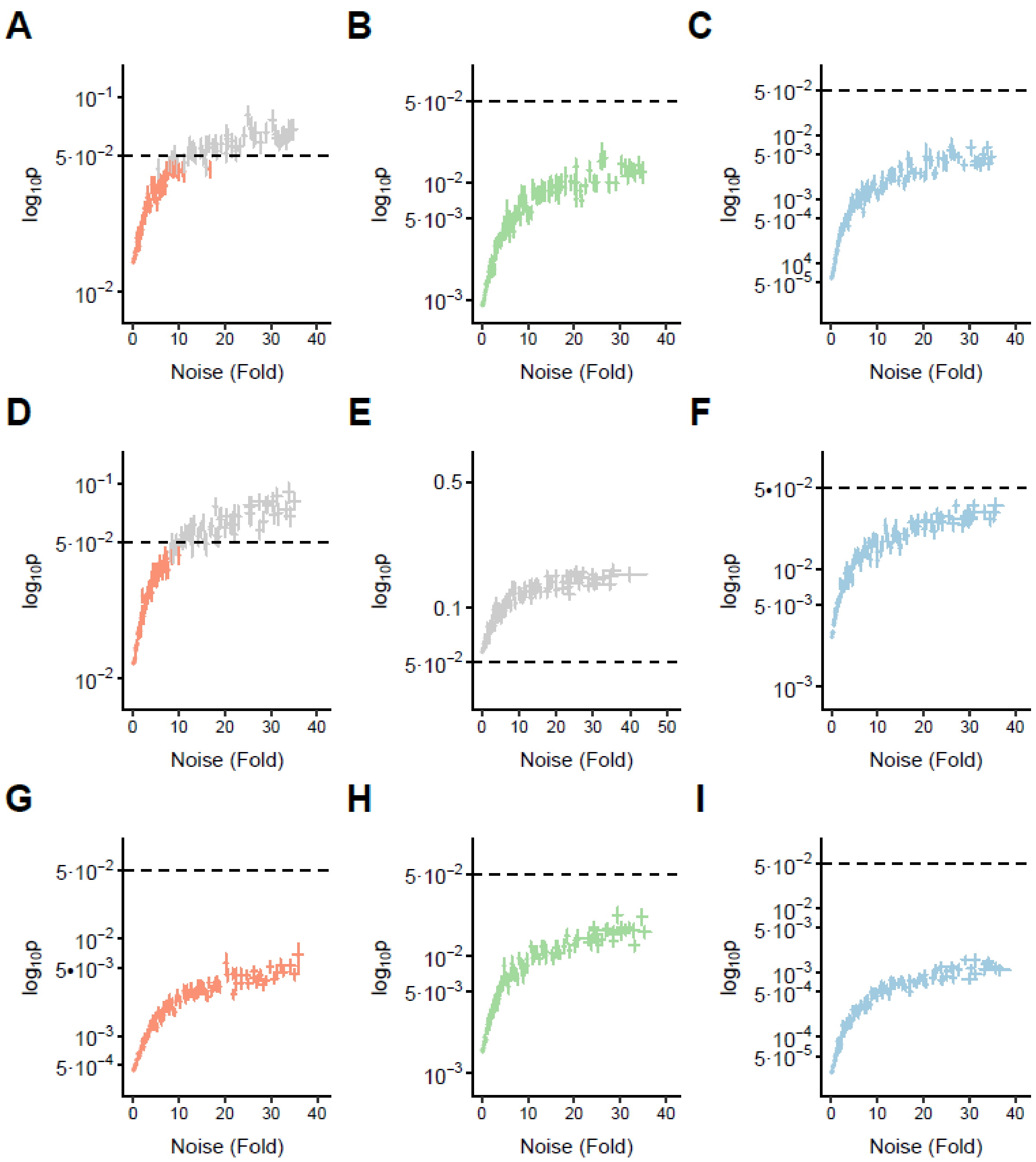


**Figure S4. P-values for supersaturation estimates for aggregating proteins in individual hereditary protein myopathies are robust against random noise.** Random noise from increasingly wide Gaussian distributions was introduced into protein supersaturation scores for the proteome and proteins enriched in affected fibers from hereditary myopathies, as shown in **Figure 2**. Points plotted are the mean ± S.E.M. of one-tailed Wilcoxon/Mann-Whitney p-values between aggregate and proteome from 100 trials at each noise level based on: healthy control (**a-c**), disease control (**d-f**), or affected fiber context (**g-i**) for desminopathy (**a**, **d**, **g**), filaminopathy (**b**, **e**, **h**), and myotilinopathy (**c**, **f**, **i**), respectively. A one-tailed one-sample Student's t-test was performed at each noise level to determine whether p-values were significantly less than 0.05 **(d-f)**. Colored points represent significant results and grey points represent non-significant results. Dashed line marks p=0.05.

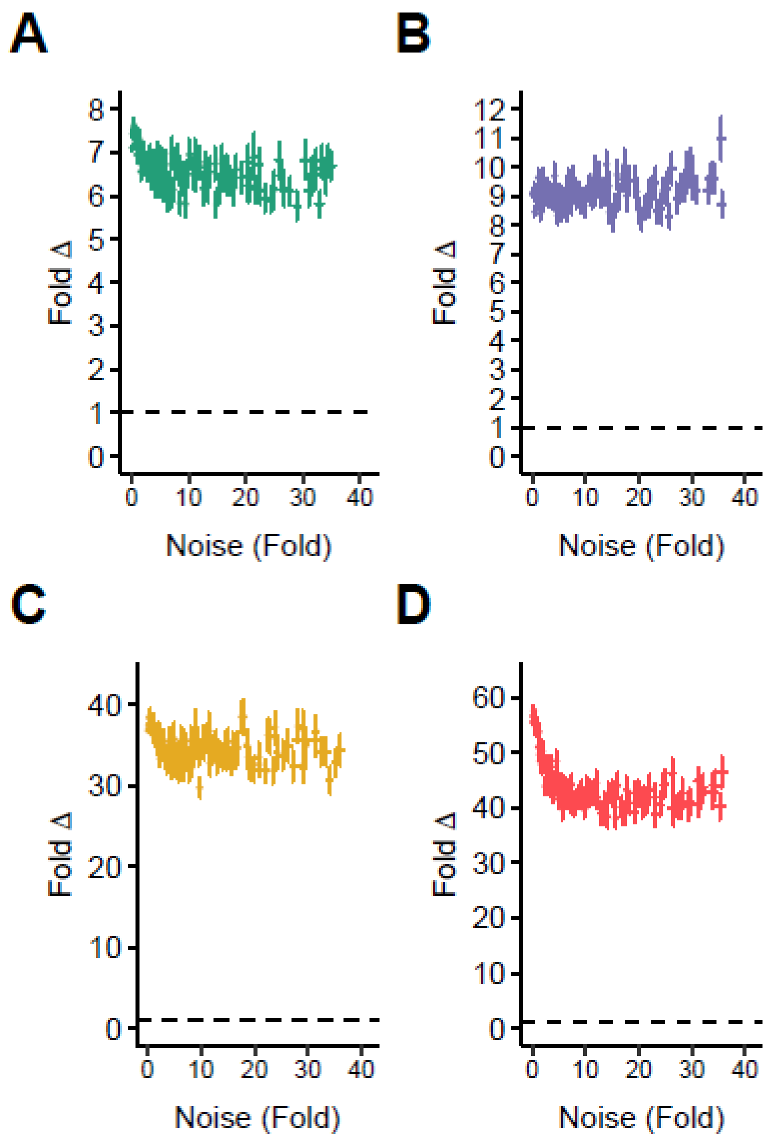


**Figure S5. Fold change for supersaturation estimates for IBM RV-enriched proteins are robust against random noise.** Random noise from increasingly wide Gaussian distributions was introduced into proteome and RV-enriched protein supersaturation scores shown in **Figure 3**. Points plotted are the mean ± S.E.M. of median fold difference between aggregate and proteome from 100 trials at each noise level based on: **(a)** healthy control, (**b**) disease control, (**c**) affected fiber, or (**d**) rimmed vacuole context. A one-tailed one-sample Student's t-test was performed at each noise level to determine whether median fold differences were significantly greater than 1 **(d-f)**. Colored points represent significant results. Dashed line marks median fold difference of 1.

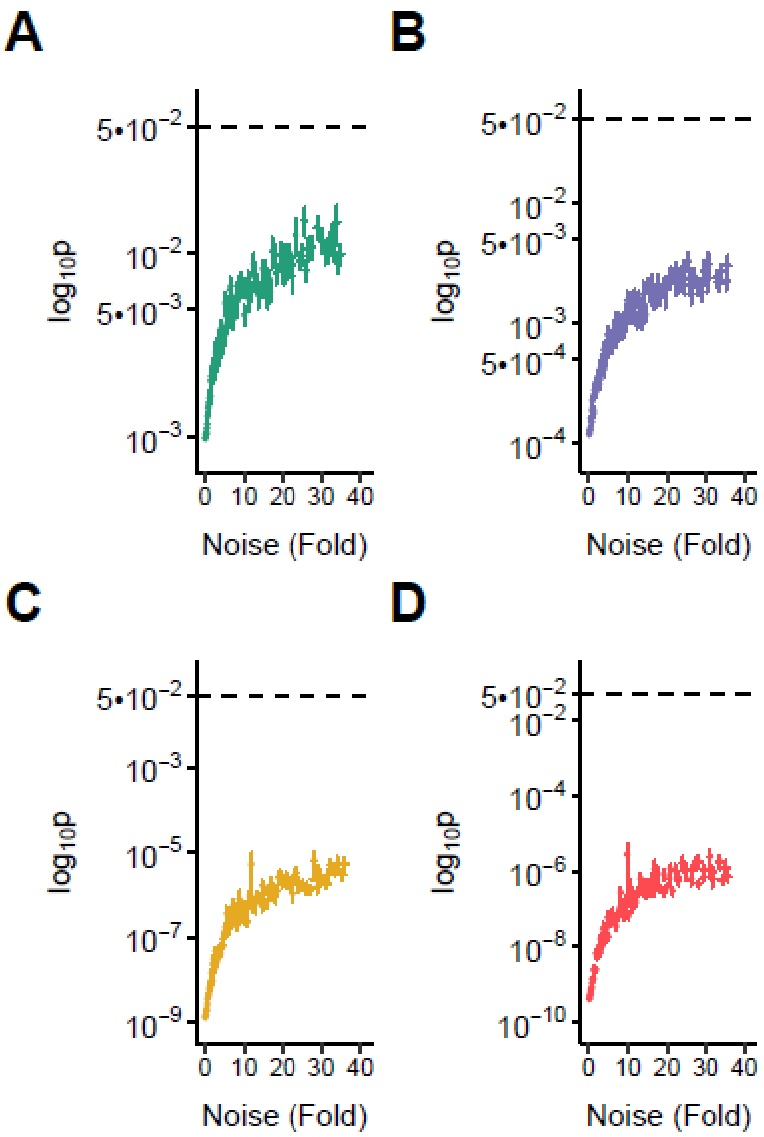


**Figure S6. P-values for supersaturation estimates for IBM RV-enriched proteins are robust against random noise.** Random noise from increasingly wide Gaussian distributions was introduced into proteome and aggregate protein supersaturation scores shown in **Figure 2**. Points plotted are the mean ± S.E.M. of one-tailed Wilcoxon/Mann-Whitney p-values between aggregate and proteome from 100 trials at each noise level based on: **(a)** healthy control, (**b**) disease control, (**c**) affected fiber, or (**d**) rimmed vacuole context. A one-tailed one-sample Student's t-test was performed at each noise level to determine whether p-values were significantly less than 0.05 **(d-f)**. Colored points represent significant results and grey points represent non-significant results. Dashed line marks p=0.05.

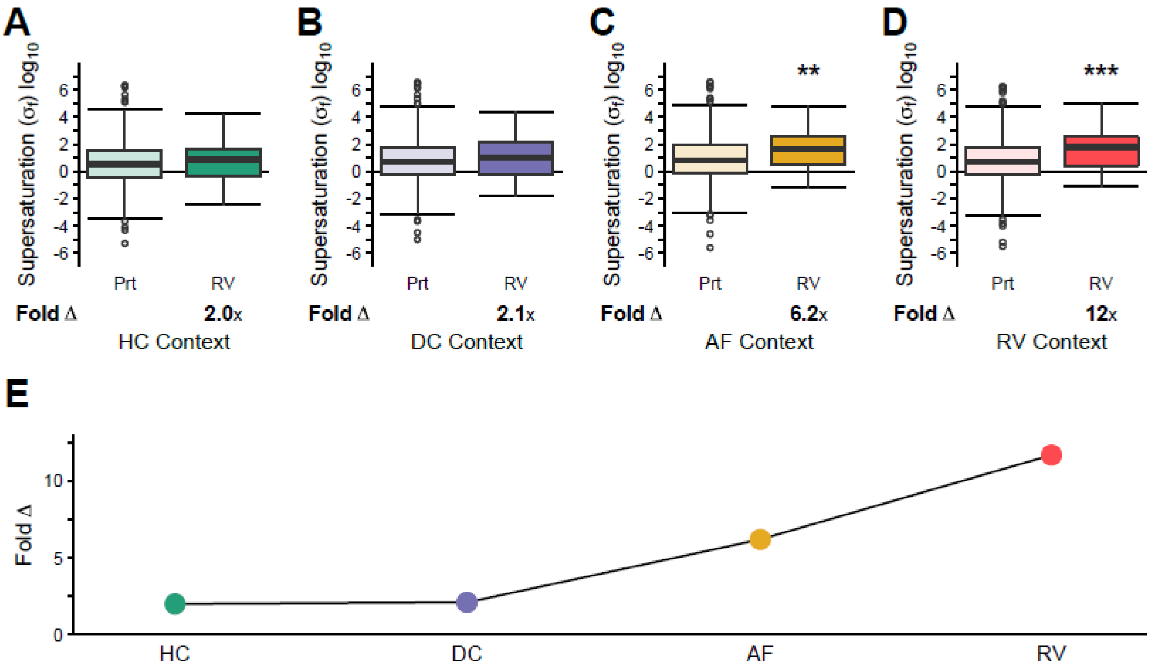


**Figure S7. Escalating supersaturation in inclusion body myositis for proteins with coverage across sample types.** Comparison of supersaturation scores ($\sigma_f$) for the proteome (Prt N=830) and proteins enriched in rimmed vacuoles (RV N=47) relative to diseased control myofibers. In this analysis, only proteins detected in all IBM sample types (healthy control myofibers (HC), control myofibers unaffected in diseased samples (DC), aggregate-containing affected myofibers (AF), and rimmed vacuoles (RV)) are included. Supersaturation scores for: **(a)** HC, **(b)** DC, **(c)** AF, and **(d)** RV. **(e)** Comparison of the fold difference in median $\sigma_f$ between RV and Prt. Box plots and statistical tests as in **Figure 1**. $^{**}p < 0.01$, $^{***}p < 0.001$.

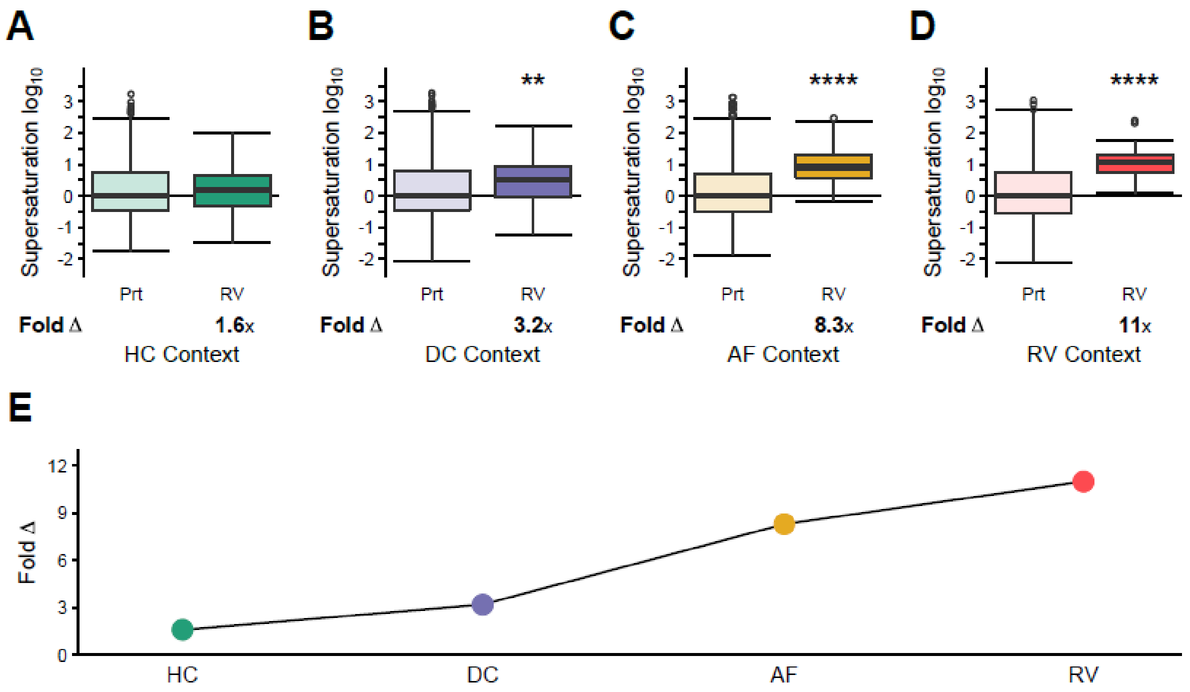


**Figure S8. Escalating supersaturation in inclusion body myositis using $Z_{agg}$.** Comparison of supersaturation scores derived from protein abundances and $Z_{agg}$ for the proteome (Prt) and proteins enriched in rimmed vacuoles (RV) relative to diseased control myofibers. **(a)** Healthy control myofiber (HC) (Prt N=1534, RV N=45), **(b)** control myofibers unaffected in diseased samples (DC) (Prt N=1883, RV N=50), **(c)** aggregate-containing affected myofibers (AF) (Prt N=2263, RV N=50), and **(d)** rimmed vacuoles (RV) (Prt N=2025, RV N=50). **(e)** Comparison of the fold difference in median $\sigma$ between RV and Prt. Box plots and statistical tests as in **Figure 1**. $^{**}p < 0.01$, $^{****}p < 0.0001$.

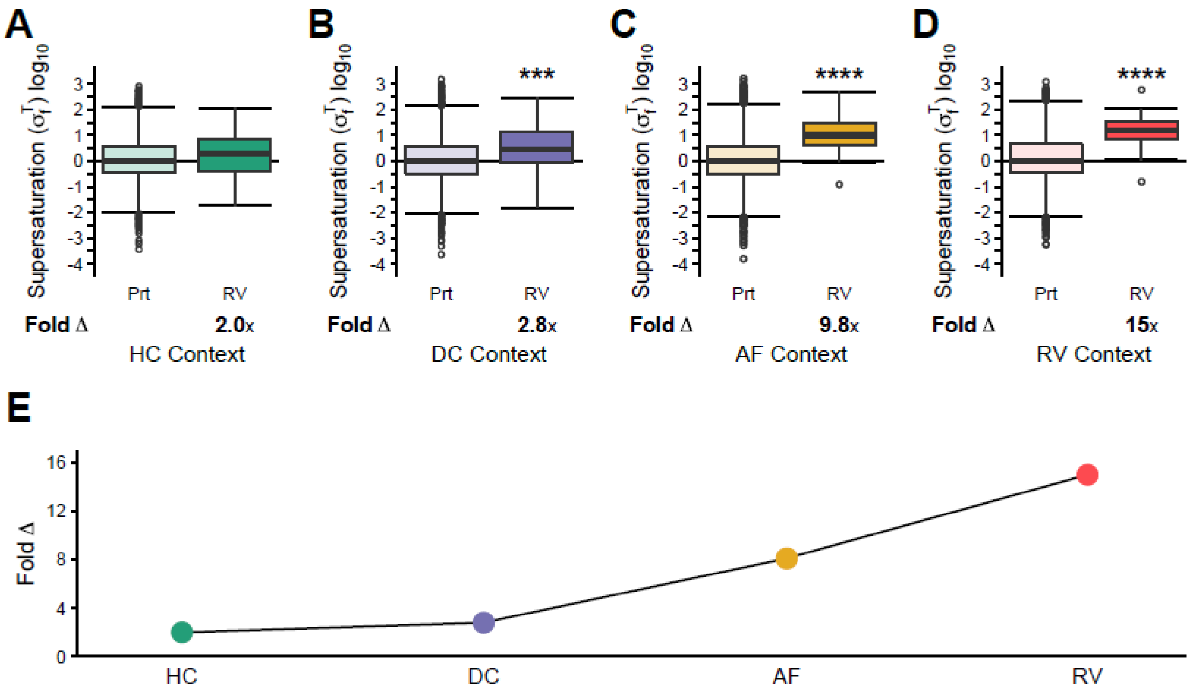


**Figure S9. Escalating supersaturation in inclusion body myositis using TANGO.** Comparison of supersaturation scores ($\sigma_f^T$) derived from protein abundances and TANGO for the proteome (Prt) and proteins enriched in rimmed vacuoles (RV) relative to diseased control myofibers. **(a)** Healthy control myofiber (HC) (Prt N=1646, RV N=48), **(b)** control myofibers unaffected in diseased samples (DC) (Prt N=2050, RV N=53), **(c)** aggregate-containing affected myofibers (AF) (Prt N=2445, RV N=53), and **(d)** rimmed vacuoles (RV) (Prt N=2165, RV N=53). **(e)** Comparison of the fold difference in median $\sigma_f^T$ between RV and Prt. Box plots and statistical tests as in **Figure 1**. ***$p < 0.001$, ****$p < 0.0001$.

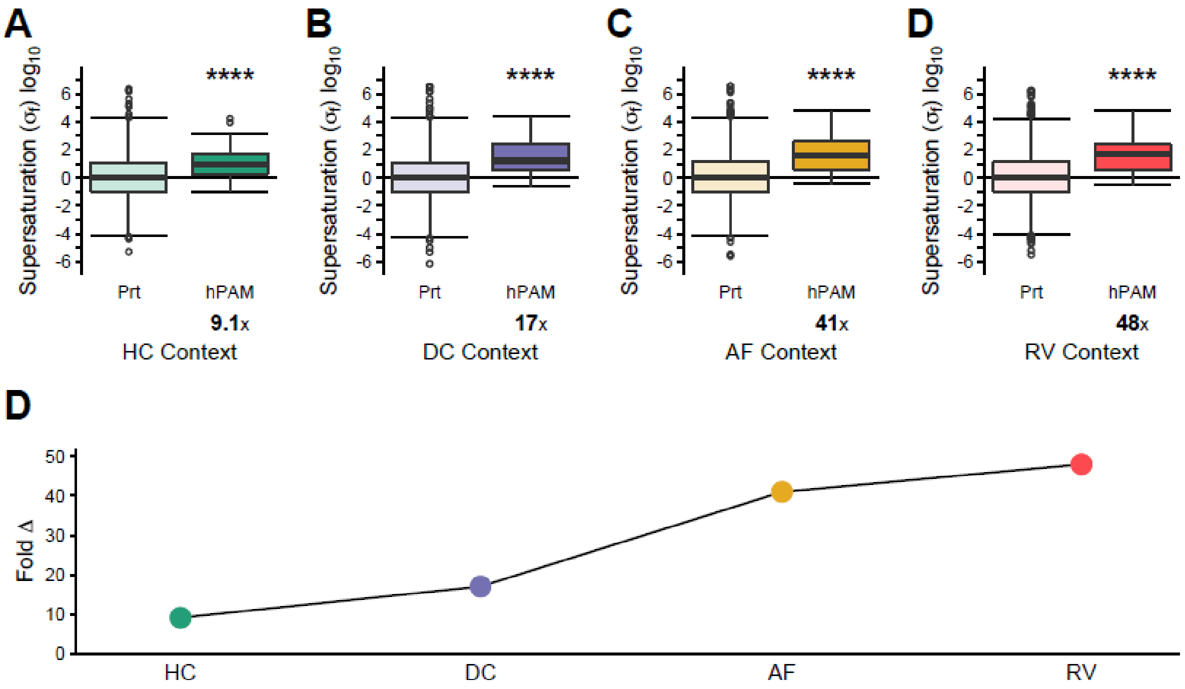


**Figure S10. Escalating supersaturation for hPAM aggregate-enriched proteins in the sporadic context.** Comparison of supersaturation scores ($\sigma_f$) calculated based on IBM context for the proteome (Prt) and proteins enriched in aggregates from hPAMs relative to diseased control myofibers (hPAM). Supersaturation scores and protein abundances for: **(a)** healthy control myofiber (HC) (Prt N=1605, hPAM N=46), **(b)** control myofibers unaffected in diseased samples (DC) (Prt N=1988, RV N=50), **(c)** aggregate-containing affected myofibers (AF) (Prt N=2396, RV N=50), and **(d)** rimmed vacuoles (RV) (Prt N=2104, RV N=50). **(e)** Comparison of the fold difference in median $\sigma_f$ between hPAM and Prt. Box plots and statistical tests as in **Figure 1**. ****$p < 0.0001$.

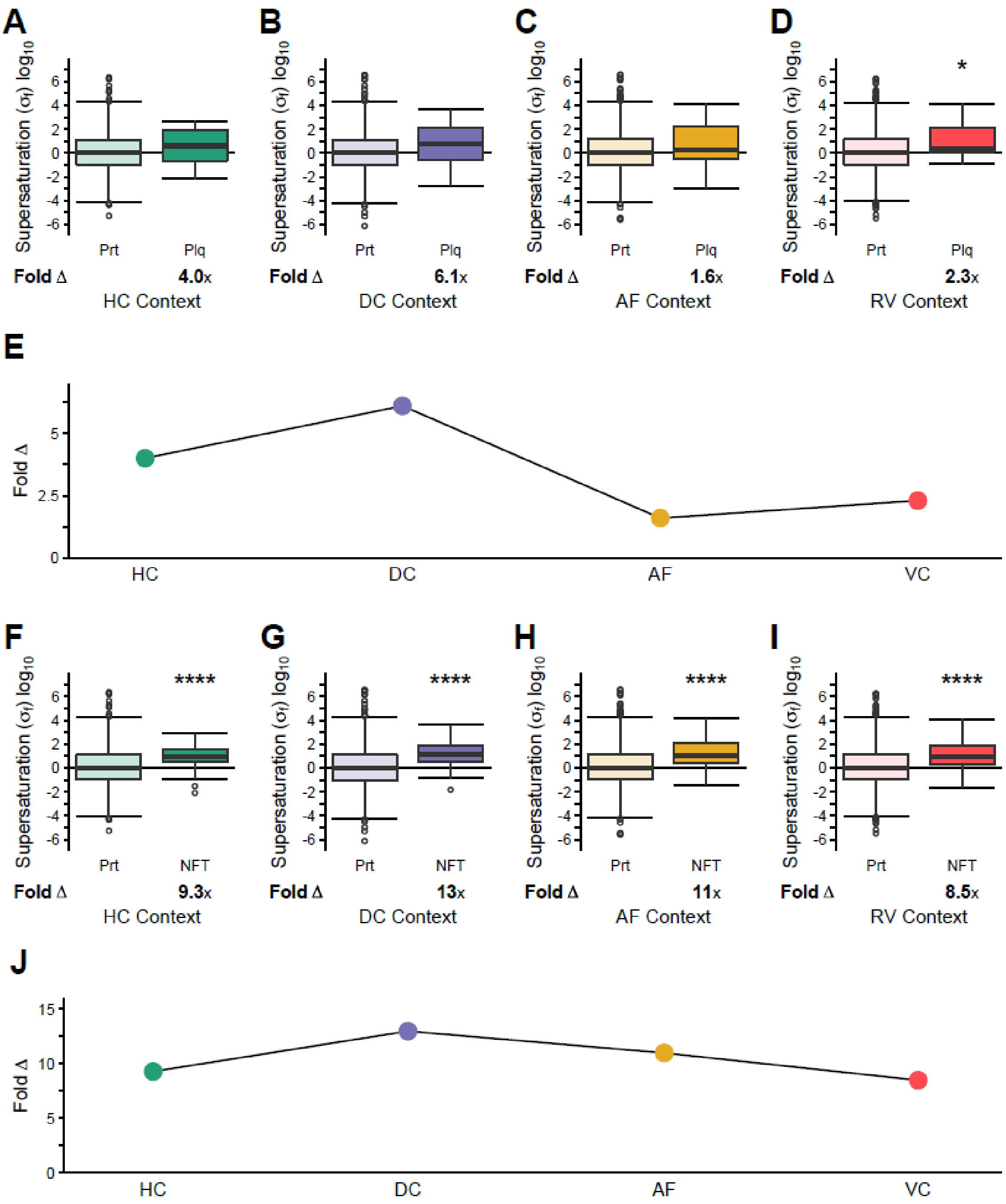


**Figure S11. Plaque- and NFT-enriched proteins do not exhibit escalating supersaturation scores in IBM tissues.** Comparison of supersaturation scores ($\sigma_f$) for the proteome (Prt) and proteins enriched in plaques (Plq, **a-d**) or neurofibrillary tangles (NFT, **f-i**). Supersaturation scores and protein abundances for: **(a, f)** healthy control myofiber (HC) (Prt N=1605, Plq N=16, NFT N=41), (**b, g)** control myofibers unaffected in diseased samples (DC) (Prt N=1988, Plq N=16, NFT N=42), **(c, h)** aggregate containing affected myofibers (AF) (Prt N=2396, Plq N=20, NFT N=48), and **(d, i)** rimmed vacuoles (RV) (Prt N=2104, Plq N=18, NFT N=48). Fold difference in median $\sigma_f$ between proteome and **(e)** Plq or **(j)** NFT. Box plots and statistical tests as in **Figure 1**. *p < 0.05, ****p < 0.0001.

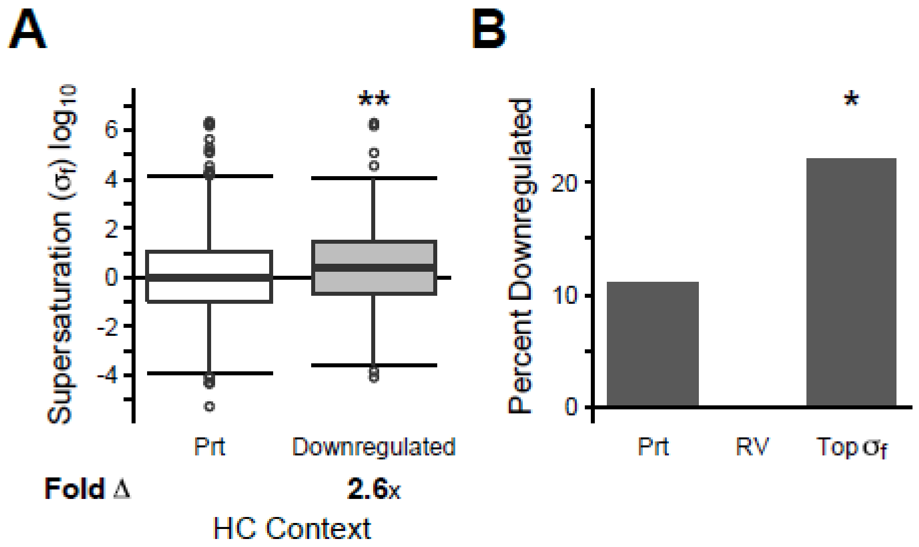


**Figure S12. Protein supersaturation is associated with downregulation utilizing RNAseq datasets.** (**a,b**) Only proteins with defined $\sigma_f$ scores in HC for which transcripts were detected in both Ctl and sIBM RNA sequencing data are included. **(a)** Supersaturation scores ($\sigma_f$) for the proteome (Prt) (N=1366) and proteins downregulated from Ctl to sIBM RNA sequencing data (N=157). Box plots and statistical tests as in **Figure 1**. **(b)** Percentage of proteins downregulated in the proteome (Prt) (157/1366), amongst proteins enriched in rimmed vacuoles (RV) (0/42), and amongst the top 5% most supersaturated proteins (based on HC context) (Top $\sigma_f$) (15/68).

**Dataset S1. Proteins enriched in rimmed vacuoles.** List of proteins enriched in rimmed vacuoles relative to unaffected fibers from diseased samples, with annotation as to whether they have previously been identified in IBM muscle, in what type of aggregate structure they have been found, and whether they have an associated mutation known to cause myopathy.

**Dataset S2. Proteins enriched in plaques, neurofibrillary tangles, and protein aggregation myopathies.** List of proteins enriched in plaques and neurofibrillary tangles or co-aggregating with TDP-43 are based on previously reported data (12-14). Proteins enriched in affected fibers from desminopathy, filaminopathy, and myotilinopathy also shown.

**Dataset S3. Aggregation propensity scores**. $Z_{agg}$, $Z_{agg}^{SC}$, and TANGO scores (15) calculated as described in Methods..

**Dataset S4. mRNA expression levels.** Cross-tissue mRNA, skeletal muscle mRNA microarray, control skeletal muscle RNA sequencing (FKPM), and sIBM RNA sequencing (FKPM) expression data.

**Dataset S5. Hereditary protein aggregation myopathy abundance data.** Protein abundance based on mass spectrometry data for desminopathy, filaminopathy, and myotilinopathy, collected and calculated as described in Methods.

**Dataset S6. Sporadic inclusion body myositis abundance data.** Protein abundance based on mass spectrometry data for sporadic inclusion body myositis, collected and calculated as described in Methods.

**Dataset S7. Unfolded supersaturation scores.** Supersaturation calculated using cross-tissue mRNA expression and $Z_{agg}$ (15) ($\sigma_u$), mRNA expression from skeletal muscle microarray and $Z_{agg}$ ($\sigma_u^{ts}$) and cross-tissue mRNA expression and TANGO ($\sigma_u^T$) as described in Methods.

**Dataset S8. Hereditary protein aggregation myopathy supersaturation scores ($\sigma_f$).** $\sigma_f$ based on mass spectrometry data and structurally-corrected aggregation propensity scores ($Z_{agg}^{SC}$) for desminopathy, filaminopathy, and myotillinopathy, calculated as described in Methods.

**Dataset S9. Sporadic inclusion body myositis supersaturation scores ($\sigma_f$).** $\sigma_f$ based on mass spectrometry data and structurally-corrected aggregation propensity scores ($Z_{agg}^{SC}$) for sporadic inclusion body myositis, calculated as described in Methods.

**Dataset S10. Sporadic inclusion body myositis supersaturation scores ($\sigma_f^T$).** $\sigma_f^T$ based on mass spectrometry data and TANGO scores for sporadic inclusion body myositis, calculated as described in Methods.

**Dataset S11. Upregulated and downregulated proteins in sporadic inclusion body myositis.** Proteins upregulated and downregulated in sporadic inclusion body myositis affected fibers compared to healthy controls based on proteomic and RNA sequencing data.

**Dataset S12. Summary of statistical analysis and families of statistical tests.** Individual families of tests for the purposes of multiple hypothesis correction are shown under each bold heading, with corresponding figure to which the results are relevant. P-values are shown, with the method used to obtain the p-value, and the starred significance based on $*p < 0.05$, $**p < 0.01$, $***p < 0.001$, $****p < 0.0001$.
